# Organizational Hierarchy and Structural Diversity of Microvascular Pericytes in Adult Mouse Cortex

**DOI:** 10.1101/114777

**Authors:** Roger I. Grant, David A. Hartmann, Robert G. Underly, Andrée-Anne Berthiaume, Narayan R. Bhat, Andy Y. Shih

## Abstract

Smooth muscle cells and pericytes, together called mural cells, coordinate many distinct vascular functions. Smooth muscle cells are ring-shaped and cover arterioles with circumferential processes, whereas pericytes extend thin processes that run longitudinally along capillaries. In between these canonical mural cell types are cells with mixed phenotype of both smooth muscle cells and pericytes. Recent studies suggest that these transitional cells are critical for controlling blood flow to the capillary bed during health and disease, but there remains confusion on how to identify them and where they are located in the brain microvasculature. To address this issue, we measured the morphology, vascular territory, and α-smooth muscle actin content of structurally diverse mural cells in adult mouse cortex. We first imaged intact 3-D vascular networks to establish the locations of major gradations in mural cell appearance as arterioles branched into capillaries. We then imaged individual mural cells occupying the regions within these gradations. This revealed two transitional cells that were often similar in appearance, but with sharply contrasting levels of α-smooth muscle actin. Our findings highlight the diversity of mural cell morphologies in brain microvasculature, and provide guidance for identification and categorization of mural cell types.

## INTRODUCTION

Arteries and arterioles are covered by smooth muscle cells, which are short, ring-shaped and densely packed. In contrast, capillaries are covered by pericytes with intermittent, protruding cell bodies and thin processes that run longitudinally along capillaries.^1^ Despite the stark contrast in appearance and vascular territory between smooth muscle cells and pericytes, these mural cells form a seamless network along the entire vascular bed. Following early description of pericytes by Rouget^2^, studies by Zimmermann^3^ and Mayer^4^ showed that smooth muscle cells and pericytes are bridged by mural cells with features of both cell types, which were referred to as “transitional forms of pericytes” or “pre-capillary pericytes”.^5^ Zimmermann emphasized the difficulty in studying different pericyte subtypes with methods of his time, which did not allow individual mural cells to be clearly visualized. Today, surveys of pericyte organization and diversity remain challenging because cellular morphology and 3-dimensional organization of blood vessels are difficult to gather from immunostained, thin tissue sections. Further, accepted immunohistochemical stains for pericytes, such as PDGFRβ and CD13 (aminopeptidase N)^1, 6^, label contiguous groups of mural cells, and only the membrane of the cells, precluding the assessment of individual cell morphologies. Consequently, the difference between mural cell types as small arterioles transition to capillaries has remained poorly defined.

The role of pericytes in cerebral blood flow regulation has been a controversial issue^7^, with some concluding that pericytes regulate cerebral blood flow *in vivo*^8-11^, while others refute this claim.^12^-^14^ However, this controversy can be partially attributed to a lack of consensus on how to define transitional forms of pericytes. For example, most groups agree that the proximal branches of penetrating arterioles are contractile *in vivo*^8, 12, 13^ The mural cells that cover these vessels appear to fit the concept of transitional pericytes, *i.e.,* a mixed phenotype with protruding cell bodies and an elongated shape like pericytes, and processes encircling the lumen like smooth muscle cells. Yet, these cells are often categorized as either smooth muscle cells^13^ or pericytes^8^ by different research groups, which inevitably leads to uncertainty as to which mural cells were studied. A more detailed characterization of the morphology of different mural cell types and the microvascular territories they occupy would help with the interpretation of existing and future studies on brain pericytes.

The goal of this study was to characterize the various mural cell types found in the adult mouse cerebral cortex. We determined whether transitional pericyte forms could be reliably distinguished from canonical smooth muscle cells and pericytes based on cell morphology, α-smooth muscle actin (α-SMA) content, location within the vascular bed, or a combination of these metrics. Three-dimensional cerebrovascular networks were imaged in order to examine the vascular territories occupied by different mural cell types. Further, transgenic mice with sparsely-labeled fluorescent mural cells were used to measure features of individual cells.^6^

## METHODS

The Institutional Animal Care and Use Committee at the Medical University of South Carolina approved the procedures used in this study. The University has accreditation from the Association for Assessment and Accreditation of Laboratory Animal Care International, and all experiments were performed within its guidelines. All data were analyzed and reported according to ARRIVE guidelines.

### Animals

Heterozygous male PDGFRβ-Cre^15^ mice (FVB and C57BL/6 x 129 background), a generous gift from Prof. Volkhard Lindner of the Maine Medical Center Research Institute, or NG2-CreER^TM^ mice (#008538; Jackson labs; C57BL/6 background)^16^ were bred with homozygous female Ai14 reporter mice (#007914; Jackson labs; C57BL/6 background)^17^ to produce PDGFRβ-tdTomato, and NG2-tdTomato offspring. We used both male and female offspring, between 3 to 9 months of age, for all parts of this study. As previously described^6^, PDGFRβ-tdTomato mice provided a contiguous label of nearly all mural cells throughout the cerebrovasculature, while NG2-tdTomato provided a sparse labeling of mural cells following induction of Cre recombinase expression with tamoxifen (100mg/kg i.p. dissolved 9:1 corn oil:ethanol for 1 or 2 days).

### Tissue fixation

Mice were perfusion fixed with phosphate-buffered saline (PBS), followed by 4% paraformaldehyde (PFA) in PBS through a trans-cardiac route. After perfusion, the brain was extracted and placed in 4% PFA in PBS. Brains were then transferred to PBS with 0.01% sodium azide after overnight post-fixation for longer term storage.

### Tissue processing for optically-cleared specimens

Coronal brain slices from PDGFRβ-tdTomato mice were collected at a thickness of 0.5 to 1 mm using a vibratome. Slices were first subjected to an antigen retrieval protocol. This consisted of a 1-hour incubation in a 1:2 ratio of 0.25% trypsin-EDTA (Sigma-Aldrich; T4049) and PBS at 37^o^C in a water bath. This was followed by 1-hour of washing in PBS at room temperature under slow nutation. Slices were then incubated with a FITC-conjugated α-SMA antibody (Sigma-Aldrich; F3777) for 1-week. The antibody was used at a dilution of 1:200 in an antibody solution composed of 2% Triton X-100 (v/v, Sigma-Aldrich; X100), 10% goat serum (v/v, Vector Laboratories; S1000), and 0.1% sodium azide (w/v, Sigma-Aldrich; S2002) in PBS. After 1-week of immunostaining, slices were washed in PBS for 2-hours. We then cleared the tissues using the “See Deep Brain” (SeeDB) method over 4 days.^18^ On the fifth day, slices were imaged with two-photon microscopy while immersed in full SeeDB solution. All incubations were performed at room temperature under slow nutation, with samples protected from light with aluminum foil. Negative control samples for α-SMA staining were incubated at the same time, using adjacent slices from the same animal, in solution without α-SMA antibody (**Supplementary Fig. 1**). This control confirmed that most green fluorescent signal was indeed from detection of α-SMA protein. Critically, some autofluorescence in SeeDB-cleared tissue was detected along blood vessels, including most pericyte cell bodies, which can be easily mistaken for true α-SMA staining in pericytes (**Supplementary Fig. 2**). The autofluorescence we detected in pericytes was likely not the result of spectral overlap from the tdTomato channel because the autofluorescence could be detected when exciting at 800 nm, where tdTomato excitation is inefficient.^19^ Thus, quantification of α-SMA content was not performed on SeeDB-cleared specimens.

### Tissue processing for thin sections

Brain slices from NG2-tdTomato mice were collected at thickness of 100 to 200 μm. Slices underwent the same antigen retrieval protocol mentioned above (1-hour trypsin treatment), and were then incubated overnight in antibody solution with the following additions: α-SMA primary antibody from mouse host (1:200 dilution; Sigma-Aldrich; A5228) and FITC-conjugated tomato lectin (1:250 dilution; Vector Labs; FL-1171). Following overnight incubation, we washed slices in PBS for 30 minutes, and then transferred to antibody solution containing anti-mouse Alexa 647 secondary antibody (1:500 dilution; ThermoFisher; A31626) for a 2-hour incubation period. Slices were then washed again in PBS for 30 minutes, mounted onto glass slides, and sealed with Fluoromount G (Southern Biotech; 0100-01) under a coverslip.

### Two-photon imaging of optically-cleared specimens

Imaging was performed with a Sutter Moveable Objective Microscope and a Coherent Ultra II Ti:Sapphire laser source. Cleared tissues were mounted in a small petri dish immersed in SeeDB solution, then covered with a 100 μm thick glass coverslip. Imaging was performed at 975 nm excitation under a 20-X, 1.0 NA water-immersion objective (XLUMPLFLN; Olympus). Green and red fluorescence emission was collected through 515/30 (Chroma ET605/70m-2P) and 615/60 (Chroma ET510/50m-2P) filters, respectively. Image stacks were collected in the barrel field of the primary somatosensory cortex, located by comparing brain regions with a mouse atlas.^20^ We imaged penetrating arterioles with branches that were fully contained within the tissue slice. Two to three image stacks were collected to capture the entirety of a penetrating arteriole, often spanning the pial surface to the callosum. Imaging resolution was 0.63 μm per pixel in the lateral plane (medial-lateral axis) and a z-step of 1 μm was used (anterior-posterior axis). Laser powers at 975 nm were 25 mW at the sample surface, and 220 mW at ∼400 um in depth. Image volumes were stitched using XuvTools ^21^ and viewed in Imaris software (Bitplane).

### Confocal imaging of thin sections

Imaging was performed on a Leica TCS SP2 AOBS Confocal Microscope (Leica Microsystems, Inc.) using 20-X (HC PlanAPO 20x/0.7 CS), 40-X (HCX PlanAPO CS 40x/1.25-0.75 Oil immersion), or 63-X(HCX PlanAPO CS 63x/1.4-0.6 Oil immersion) objectives, which respectively had lateral resolution of 0.73, 0.37, and 0.23 μm per pixel. Step sizes in the z-dimension were either 0.5 or 1 μm. Continuous wave lasers with 488, 543, and 633 nm excitation wavelengths were used for FITC, tdTomato, and Alexa 647, respectively. Emission was collected through a prism spectrophotometer utilizing an acousto-optical tuning filter to collect all channels simultaneously. Images were collected with dimensions of 1024 by 1024 pixels with 4-times averaging, and were viewed and analyzed in FIJI software.

### Analysis of two-photon imaging datasets

In total, we examined 52 offshoots extending from 7 penetrating arterioles, collected over 2 mice. Each penetrating arteriole offshoot emerged as a single vessel from the 0^th^ order penetrating arteriole. As the 1^st^ order branch ramified into the capillary bed, we followed each possible route with new bifurcations and recorded the observation of ovoid cell bodies, α-SMA termini, and shifts in pericyte coverage. The location of each of these features reported in **Figs. 2, 3 and Supplementary Fig. 3a** was the branch order resulting from averaging over all vascular routes for each penetrating arteriole offshoot, *i.e.* 1^st^ order branch and its downstream branches. Ovoid cell bodies were “bump on a log” somata that protruded from the wall of the vessel, a canonical feature of pericytes. The α-SMA termini were identified visually as abrupt reductions of α-SMA-FITC fluorescence. Shifts in pericyte coverage were defined as locations where pericyte processes started to appear as thin strands running longitudinally along the vessel. Inter-soma distance of pericytes was measured in 3-D data sets using the measurement points tool in Imaris software.

### Analysis of confocal imaging datasets

In NG2-tdTomato mice, we targeted data collection to four mural cell groups, decided *a priori* based on data obtained from SeeDB-cleared specimens in contiguously labeled PDGFRβ-tdTomato mice: (1) Smooth muscle cells, found on 0^th^ order penetrating arterioles with short, ring-like morphology and α-SMA staining, (2) ensheathing pericytes on proximal branches of penetrating arteriole offshoots, which covered most of the endothelium, and exhibited α-SMA staining, (3) mesh pericytes typically just downstream of the α-SMA terminus, which covered most of the endothelium but did not have α-SMA staining, and (4) thin-strand pericytes deeper in the capillary bed that possessed long, thin processes and did not have α-SMA staining. All pericyte subtypes had a clear protruding ovoid cell body, while smooth muscle cells did not. We analyzed 2-D, average-projected confocal images of individual cells. Care was taken to analyze only cells that were fully contained within the image stack. All analyses were performed in FIJI software. The intensity of α-SMA staining (Alexa 647) was measured by averaging pixel values from the far-red channel within a mask of the cell generated in the tdTomato channel and subtracting the average intensity of signal from a region of neighboring parenchyma (**Fig. 4g**). For smooth muscle cells and ensheathing pericytes, we only included cells with α-SMA intensity that was 2-fold or higher than background, because of the higher background in some α-SMA stained samples. This led to the exclusion of 3 smooth muscle cells and 3 ensheathing pericytes from the total collected. Cell length was the combined length of vasculature contacted by either the cell body or processes, measured using the ‘Segmented Line’ tool in FIJI (**Fig. 5a-d,i**). Vessel diameter was measured by manually drawing a line across the vessel width at the location of the tdTomato-positive cell body, if present, in the channel containing FITC-lectin (**Fig. 5a-d,i & Supplementary Fig. 3b-d**). Some vessels segments with a noticeable diameter gradient were measured at either end of the cell body and then averaged for better accuracy. Coverage measurements were made by thresholding average-projected tdTomato images, taking care to ensure all cellular processes were captured in the thresholded image. We set this threshold to approximately one standard deviation above each cells mean intensity value, and applied it similarly for each mural cell group. All pixels above this threshold were included as the area of tdTomato labeling. We then manually demarcated the area of vessel beneath each mural cell in the FITC-lectin channel. The area labeled with tdTomato was then divided by the vessel area to provide a measure of coverage (**Fig. 5e-h,j**).

### Statistics

Pearson’s correlation tests for data from optically-cleared specimens (PDGFRβ-tdTomato) were performed in MatLab. Analyses of confocal data (NG2-tdTomato) was done with SPSS (SPSS Statistics 24, IBM). Statistical tests used for various comparisons are stated in the corresponding figure legend. Data sets were first subjected to two tests of normality, Lilliefors and Shapiro-Wilk, followed by a Levene’s test for homogeneity of variance. In accordance with the results of the previous tests, Kruskall-Wallis test with Dunn-Bonferroni *post-hoc* test was used to test for differences between pericyte subtypes in **Figs. 4g, 5i,j** and **Supplementary Fig. 3a,b**. *A priori* power analyses for sample size were not performed in these studies. The rater was blinded to mural cell group during analysis

## RESULTS

### The mural cell continuum visualized in optically-cleared tissues

To visualize mural cells and α-SMA expression within an intact microvascular network in PDGFRβ-tdTomato mice, we immunostained and then optically cleared 0.5 to 1 mm thick coronal brain slices containing the primary barrel field of the somatosensory cortex. We collected high-resolution images using two-photon microscopy, and stitched adjacent image volumes to produce large data sets that followed penetrating arterioles as they descended from the pial surface to the corpus callosum (**Fig. 1a-c, Supplementary video 1**). Our studies focused on cortical penetrating arterioles and their offshoots in upper to mid-cortex, because most *in vivo* two-photon imaging studies of cerebral blood flow have focused on this region of the brain.^8, 13, 22^

**Figure 1.**
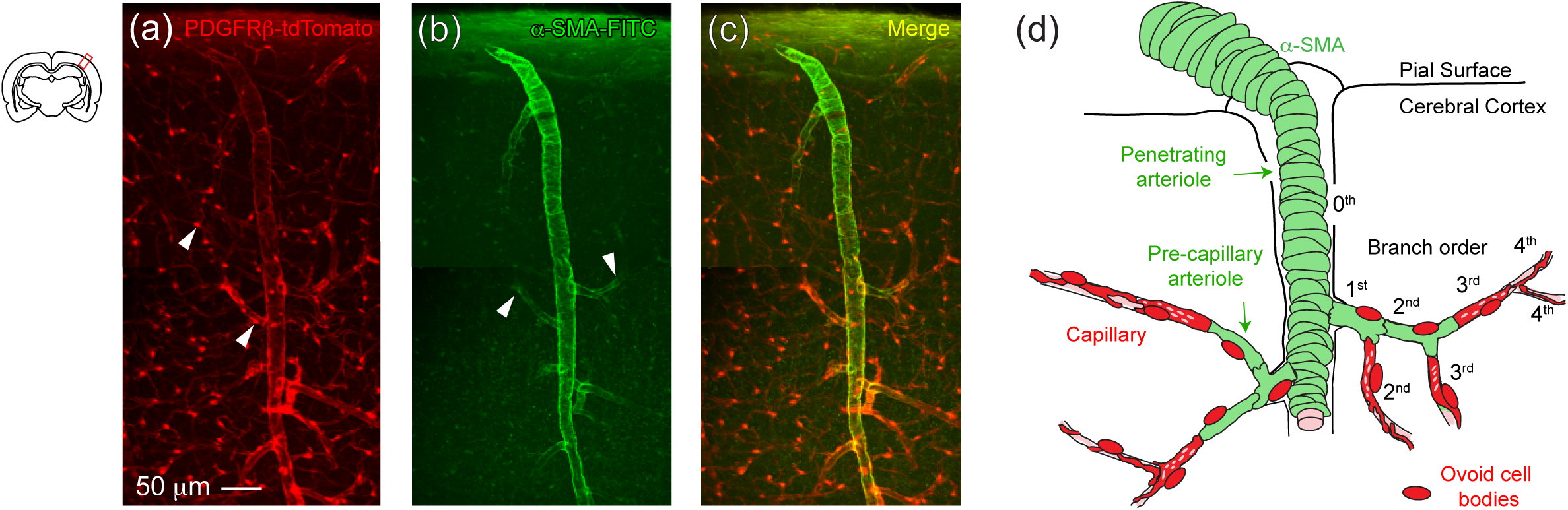
Mural cell organization and vascular structure revealed in optically-cleared mouse cortex. (**a**) Reconstructed volume from barrel cortex of a PDGFRβ-tdTomato mouse, showing tdTomato fluorescence. (**b**) The same tissues were immunolabeled with FITC-conjugated α-SMA antibody. (**c**) Composite image of tdTomato and FITC channels. (**d**) Schematic showing the terms used to describe various portions of the vascular anatomy (left side) in this study, and the system for ordering microvessel branches as they ramified from the penetrating arteriole (right side).

### Definitions

It is essential to first explain how we chose to define microvessel types and their organization. In line with previous studies of vascular topology^8, 13^, we referred to the main penetrating arteriole trunk as the 0^th^ branch order (**Fig. 1d**; right side). The first segment of an offshoot from the penetrating arteriole trunk was called the 1^st^ order branch. Each subsequent bifurcation encountered, regardless of branch size, then increased branch order by 1. This branch ordering system, however, does not apply to the base of the penetrating arteriole, where the 0^th^ order vessel tapers and splits into multiple branches, where it becomes difficult to define the 0^th^ order vessel (**Supplementary Fig. 4**).

In addition, we used the term “pre-capillary arteriole” for any branch, 1^st^ order or greater, with α-SMA staining (**Fig. 1d**; left side). Microvessels beyond the pre-capillary arterioles that did not express α-SMA, and were not ascending venules, were called capillaries. While currently a controversial issue^7^, we chose to refer to all mural cells residing on both pre-capillary arterioles and capillaries as various forms of pericyte. This is because we found that these cells possessed well-delineated, protruding cell bodies (*i.e.,* “bump on a log”)-a classic pericyte feature described by Zimmermann^3^ and others.^1, 23^ Historically, α-SMA has also been described to be expressed by a subset of brain pericytes residing on pre-capillary arterioles.^24^-^26^ However, it should be noted that there is also a different view, where all α-SMA-positive cells on a penetrating arteriole off-shoot have been referred to as pre-capillary smooth muscle cells (see Discussion).^7, 13^ Critically, α-SMA content should not be construed as a means to identify pericytes with or without contractile ability. Rather, it is included because α-SMA content is often described in pericyte studies, and the monoclonal antibody for this protein provides high specificity and a high signal to noise ratio.

### Pericyte transitions along penetrating arteriole offshoots

We focused specifically on larger penetrating arterioles that extended to cortical layers 5 and 6, rather than smaller penetrating arterioles terminating at or before layer 4.^27^ In total, 52 penetrating arteriole offshoots (1^st^ order branch and associated downstream branches) were examined over 7 penetrating arterioles from 2 mice. For each offshoot, we followed all daughter branches and recorded the branch orders at which each of the following first emerged: (1) a protruding ovoid cell body (**Fig. 2a**; arrowhead), (2) an abrupt cutoff in α-SMA staining, which we called an “α-SMA terminus” (**Fig. 2b**; arrow), and (3) a shift in pericyte coverage, a qualitative measure of where pericyte processes transitioned to a string-like appearance (**Fig. 2c**; arrowhead). We report the average branch order of the occurrence of these pericyte features for each offshoot examined.

**Figure 2.**
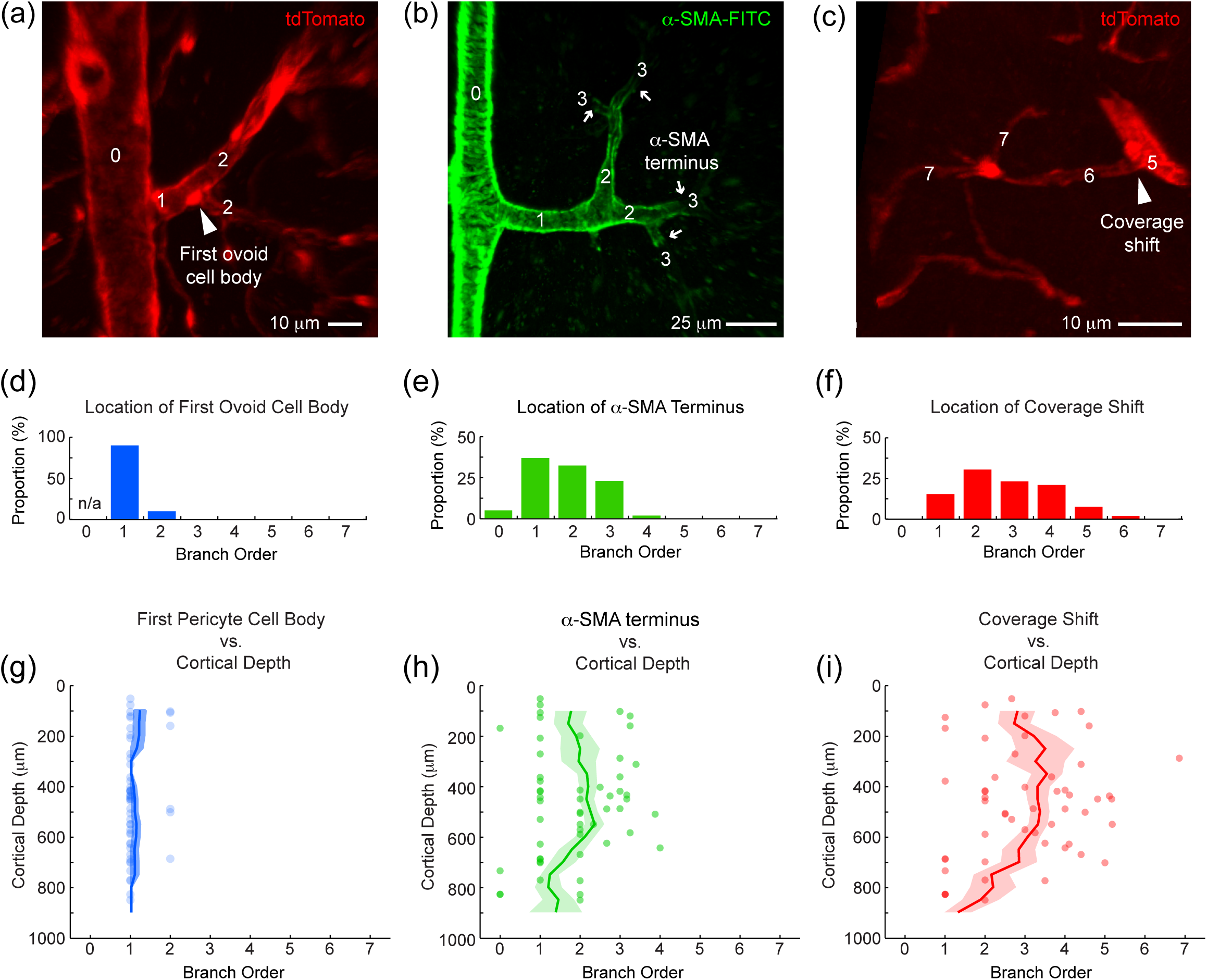
Minimal variation of pericyte features with penetrating arteriole offshoots at different cortical depths. Penetrating arterioles imaged in PDGFRβ-tdTomato mouse cortex. (**a**) The first ovoid cell body on a penetrating arteriole offshoot (arrowhead). The numbers denote branch order, as defined in Figure 1d. (**b**) Examples of α-SMA termini, where α-SMA labeling decreases sharply (arrows). (**c**) Example of a coverage shift (arrowhead), where thin, longitudinal running processes are first observed as one moves from pre-capillary arteriole to capillary. (**d-f**) Histograms showing the frequency at which each mural cell feature occurs at each branch order. (**g-i**) Scatterplots of each mural cell feature, showing branch order of occurrence as a function of cortical depth. Running average (window size 200 μm; step size 50 μm) +/-SEM is shown.

We found that ovoid cell bodies emerged on 1^st^ order branches for nearly all offshoots inspected (**Fig. 2d**); 10% first occurred on 2^nd^ order branches, but were on or immediately after the bifurcation of the 1^st^ and 2^nd^ order branches. We did not find ovoid cell bodies on the penetrating arteriole itself (0^th^ order) in upper to mid-cortex. The location of α-SMA termini ranged from 0^th^ to 4^th^ order branches (**Fig. 2e**), with 1^st^ and 3^rd^ order being the most common locations. In this case, a 0^th^ order terminus referred to a branch where α-SMA expression never extended beyond the penetrating arteriole trunk, but the trunk itself was labeled with α-SMA. Finally, shift in pericyte coverage was observed over a broad range of branch orders, from 1^st^ to 6^th^ (**Fig. 2f**). While there was a relatively high variance for the location of α-SMA termini and shift in pericyte coverage among offshoots, the average branch order of all features examined was fairly uniform over different cortical depths (**Fig. 2g-i**). For shift in pericyte coverage, a shallow peak toward higher average branch order was observed at ∼400 μm in cortical depth, which roughly corresponds to layer 4 of the cortex (**Fig. 2i**).^27^

### α-Smooth muscle actin and pericyte coverage extends farther with larger branches

We noticed that penetrating arteriole offshoots with small diameter sometimes had no detectable α-SMA (**Fig. 3a**), while branches with larger diameters could support α-SMA expression up to 4^th^ order (**Fig. 3b**). We therefore asked how the location of first ovoid cell bodies, α-SMA termini and pericyte coverage shift related to the diameter of the 1^st^ order branch (**Fig. 3a,b**; distance between arrowheads). While no correlation was found between location of first ovoid cell bodies and branch diameter (**Fig. 3c**), the locations of both α-SMA termini and coverage shifts were strongly correlated with 1^st^ order branch diameter (**Fig. 3d,e**). Thus, on larger penetrating arteriole offshoots, a step-wise transition of pericyte features could be observed: Ovoid cell bodies emerge (∼1^st^ order) → α-SMA expression terminates (∼3^rd^ to 4^th^ order) → vessel coverage shifts (∼5^th^ to 6^th^ order). These transitions occurred more quickly on smaller penetrating arteriole offshoots.

**Figure 3.**
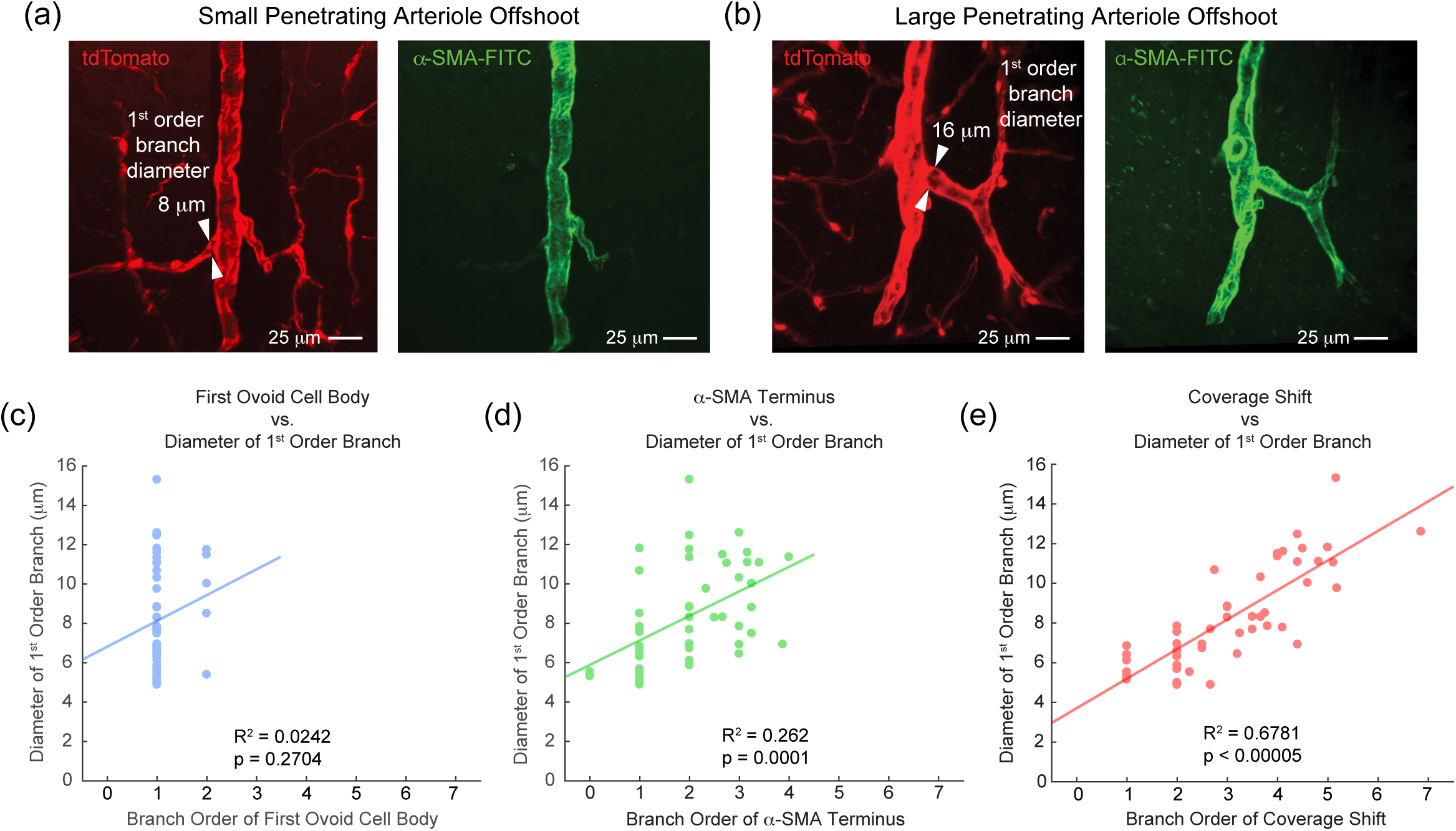
α-SMA content and pericyte coverage extend further along larger penetrating arteriole offshoots. Penetrating arterioles imaged in PDGFRβ-tdTomato mouse cortex. First order branches of penetrating arteriole offshoots range in diameter. Examples of small (**a**) and large (**b**) branches, with diameters of 8 μm and 16 μm at their points of emergence, respectively. Note that α-SMA does not extend into the small branch, while the large branch supports α-SMA for several branch orders. (**c-e**) Scatterplots of each mural cell feature, showing average branch order of occurrence as a function of the 1^st^ order branch diameter. Analysis was performed with Pearson’s correlation; n = 52 penetrating arteriole offshoots, collected over 7 penetrating arterioles from 2 mice.

### Heterogeneity of pericyte characteristics

We suspected that the step-wise transition of pericyte features in PDGFRβ-tdTomato mice reflected the presence of distinguishable pericyte types. Unlike PDGFRβ-tdTomato mice, where labeling of mural cells was continuous, the structure of individual mural cells could be examined in sparsely-labeled NG2-tdTomato mice receiving 1-2 days of tamoxifen. We therefore examined the characteristics of individual pericytes in NG2-tdTomato mice (**Fig. 4a,b**)^6^. While all pericytes sampled possessed ovoid cell bodies, the processes extending from pericyte cell bodies and their degree of vessel coverage were highly varied (**Fig. 4d-f**, see tdTomato channel) compared to the more uniform ring-shaped smooth muscle cells on 0^th^ order vessels (**Fig. 4c**).

**Figure 4.**
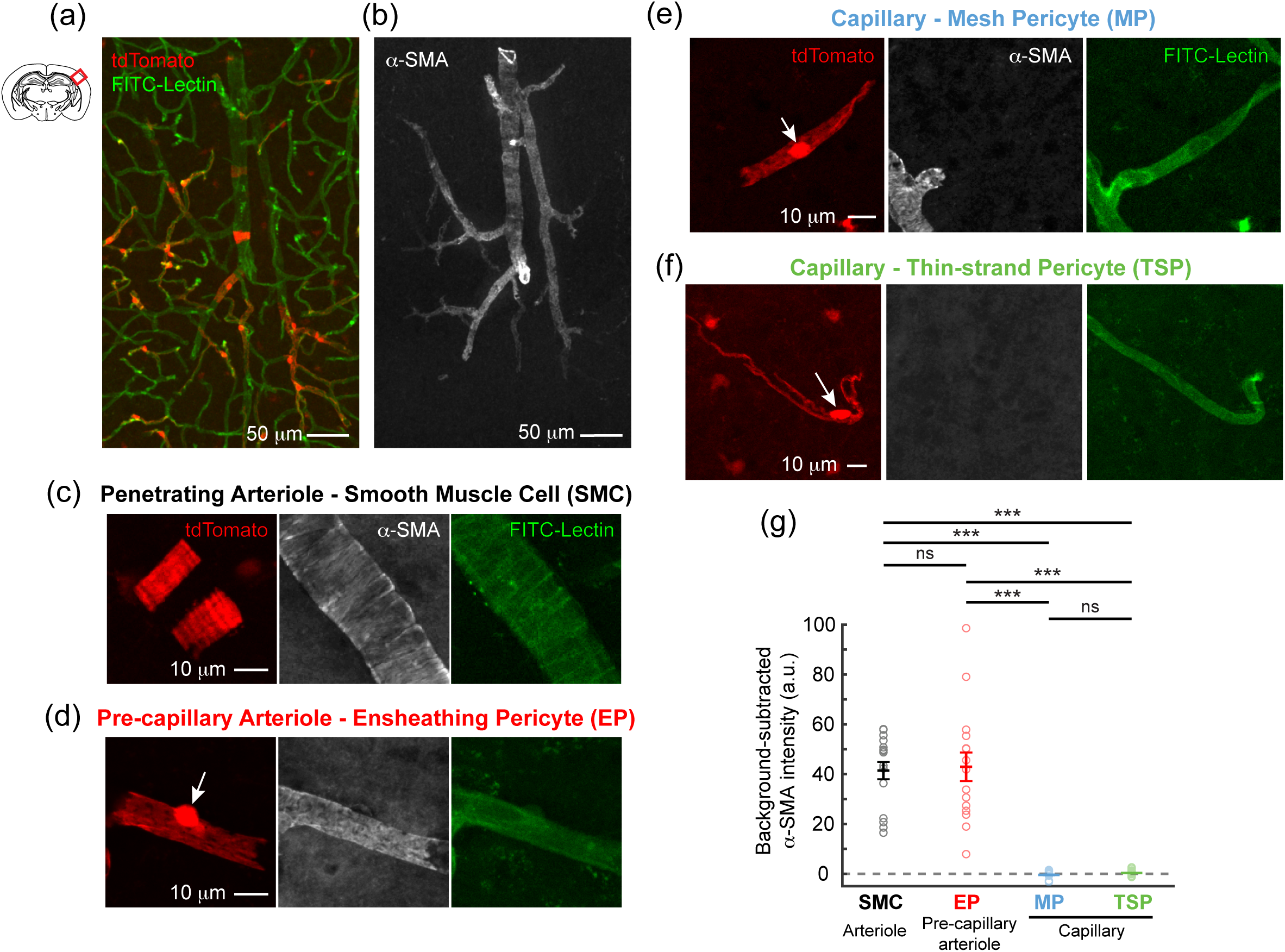
α-SMA content of mural cell types in sparsely labeled NG2-tdTomato mice. (**a**) Wide field view of penetrating arteriole in barrel cortex of NG2-tdTomato mouse. Mural cells are labeled with tdTomato (red) and vascular endothelium was labeled with FITC-conjugated lectin (green). Images were captured from 100 to 200 μm thick coronal brain sections using confocal microscopy. (**b**) The same region of tissue showing immunolabel with α-SMA antibody. (**c**) A smooth muscle cell (SMC) observed on the 0^th^ order penetrating arteriole. An α-SMA antibody and FITC-lectin co-label is also shown. (**d**) A representative ensheathing pericyte (EP) on a pre-capillary arteriole. White arrows point to ovoid cell bodies. (**e**) A representative mesh pericyte (MP) on a capillary. Note that the cell abuts the α-SMA terminus. (**f**) A typical thin-strand pericyte (TSP), the canonical form of pericyte, on a capillary. (**g**) Intensity of α-SMA exhibited for each mural cell groups. ***p<0.001, Kruskal-Wallis H test with Dunn-Bonferroni *post-hoc* test; n = 15 each for SMC, EP, MP, and TSP. Data was collected over 3 mice. Data shown as mean ± SEM. Images of α-SMA staining are raw, with no correction for background fluorescence.

We provided names for three apparent pericyte types, based upon the appearance of their processes: ensheathing, mesh and thin-strand, building upon our past descriptions from an earlier study.^6^ Ensheathing pericytes exhibited higher coverage of the vessel and were upstream of the α-SMA terminus, *i.e.,* 1^st^ to 4^th^ order branches. Mesh pericytes exhibited intermediate levels of vessel coverage, but were typically just downstream of the α-SMA terminus and therefore α-SMA-negative. Thin-strand pericytes had long thin processes that coursed along the capillary for longer distances, and were also downstream of the α-SMA terminus. In the following sections, we compared α-SMA expression, vascular territory (branch order and vessel diameter), and cellular morphology for each pericyte type and smooth muscle cells.

### α-Smooth muscle actin expression between pericyte types

We first compared the intensity of α-SMA content between mural cells (**Fig. 4d-f** and **Table 1**) The level of α-SMA immunofluorescence in ensheathing pericytes was comparable to smooth muscle cells. In contrast, α-SMA expression was undetectable in both mesh and thin-strand pericytes (**Fig. 4g**). Note how the ensheathing and mesh pericytes of **Fig. 4d,e** can be quite close in morphological resemblance but completely different in α-SMA content. We sometimes observed pericytes with faint α-SMA staining adjacent to pericytes with strong α-SMA staining (**Supplementary Fig. 5**).^6^ We decided to categorize faintly α-SMA cells as an ensheathing pericyte, in line with our definitions above (1 of 15 ensheathing pericytes in this study). Thus, α-SMA expression was used to segregate smooth muscle cells and ensheathing pericytes from mesh and thin-strand pericytes.

**Table 1.** Metrics measured for pericyte subtypes. Statistics are provided in figure legends for cell length (Fig. 5i), vessel diameter (Supplementary Fig. 3b), vessel coverage (Fig. 5j), and α-SMA intensity (Fig 4g). ^a^ n = 20 each for SMC, EP, MP and TSP. ^b^ n = 15 each for SMC, EP, MP and TSP.

| | Vessel diameter<br>($\mu\text{m}$)<sup>a</sup> | α-SMA intensity<br>(a.u.)<sup>b</sup> | Vessel Coverage<br>(%)<sup>a</sup> | Cell length<br>($\mu\text{m}$)<sup>a</sup> |
| --- | --- | --- | --- | --- |
| Smooth muscle cell<br>(SMC) | 16.2 ± 1.3 | 41.4 ± 3.9 | 95.0 ± 0.9 | 17.6 ± 1.2 |
| Ensheathing pericyte<br>(EP) | 9.0 ± 0.4 | 42.9 ± 6.1 | 95.4 ± 0.8 | 42.2 ± 2.6 |
| Mesh pericyte<br>(MP) | 6.3 ± 0.3 | -0.5 ± 0.6 | 71.6 ± 2.2 | 100 ± 7 |
| Thin-strand pericyte<br>(TSP) | 4.9 ± 0.1 | 0.3 ± 0.3 | 51.3 ± 1.3 | 149 ± 10 |

**Figure 5.**
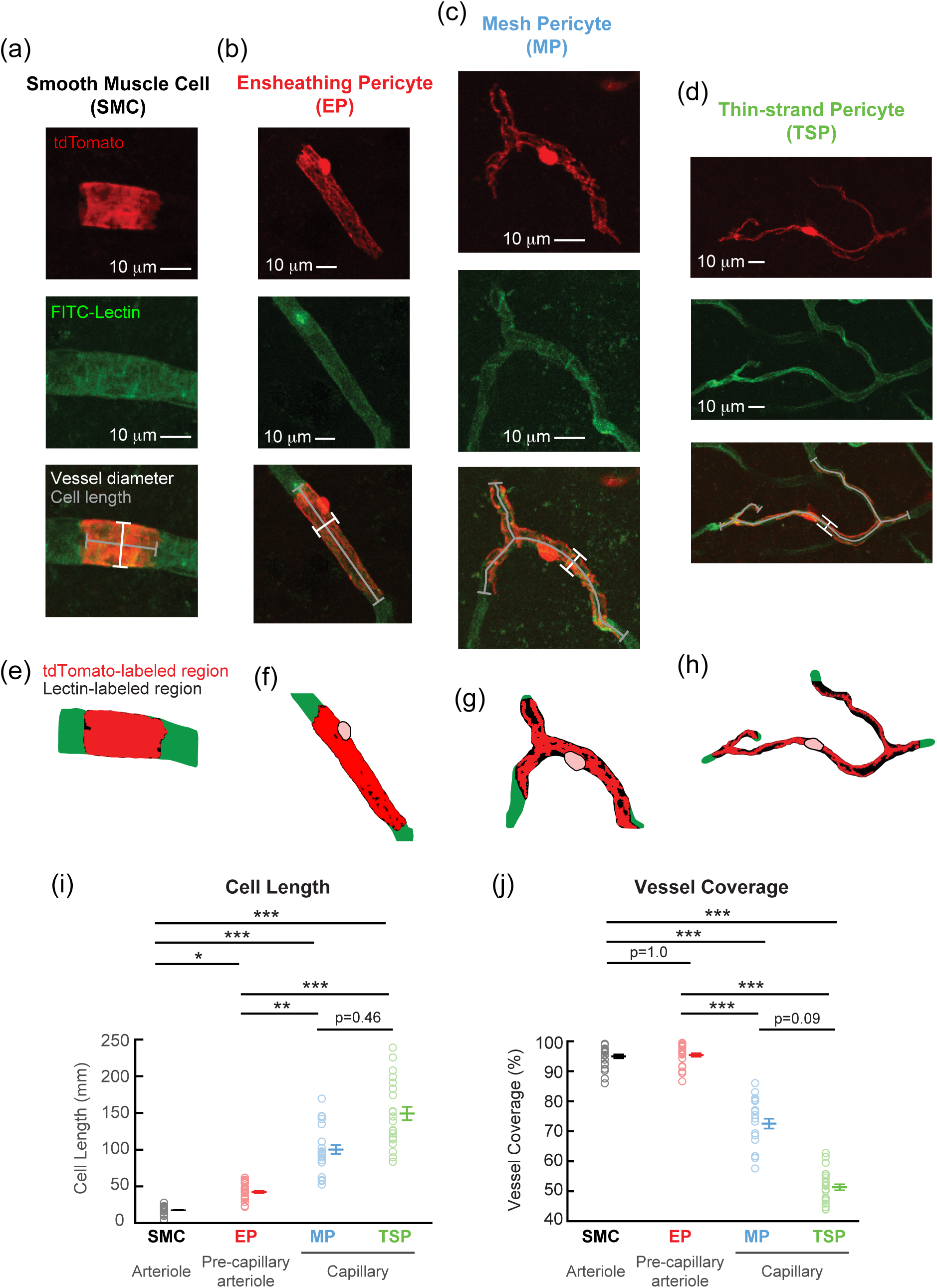
Mural cell types exhibit varying cell lengths and degrees of vessel coverage. (**a-d**) Examples of each mural cell type from NG2-tdTomato mice, with lines to measure total cell length (gray lines) and vessel diameter (white lines). (**e-h**) Vessel coverage for each mural cell was calculated by dividing tdTomato-positive area (red region) by a mask of the vessel area underlying the cell. The black regions show portions of the mask that are not covered by the mural cell. (**i,j**) Total cell length and vessel coverage for each mural cell group. *p<0.05, **p<0.01, ***p<0.001, Kruskal-Wallis H test with Dunn-Bonferroni *post-hoc* test; n = 20 each for SMC, EP, MP, and TSP. Data was collected over 3 mice. Data shown as mean ± SEM.

### Vascular territory of pericyte types

In NG2-tdTomato mice, ensheathing pericytes were found on an average branch order of 1.7 ± 0.2, mesh pericytes on 3.8 ± 0.3, and thin-strand pericytes on 5.6 ± 0.4 (**Supplementary Fig. 3a**). The branch order of ensheathing pericytes was statistically lower than thin-strand pericytes, and mesh pericytes. Mesh and thin-strand could not be distinguished based on branch order. Ensheathing pericytes were also found on microvessels with larger diameter than both mesh and thin-strand pericytes, and mesh pericytes on slightly larger microvessels than thin-strand pericytes (**Supplementary Fig. 3b and Table 1**). However, neither of these trends attained statistical significance as there was substantial overlap in the range of microvessel diameters for each pericyte type. This suggests that microvessel diameter measurements alone cannot determine the mural cell type under consideration.

### Morphological features of pericyte types

We next asked if mural cells could be statistically separated based on cell morphology. We examined cell length, where length was calculated as the total length of pericyte soma and processes in contact with FITC-lectin-labeled capillary (**Fig. 5a-d**; gray lines). As expected, smooth muscle cells on the 0^th^ order penetrating arterioles were short in length, averaging only ∼20 μm. The length of each pericyte type, however, was progressively greater than the smooth muscle cell with ensheathing, mesh and thin-strand pericytes extending over ∼40, 100, and 150 μm of vasculature, respectively (**Fig 5i and Table 1**). Ensheathing pericytes were significantly longer than smooth muscle cells, and shorter than both mesh and thin-strand pericytes, but mesh pericytes were not significantly shorter than thin-strand pericytes. Interestingly, mural cell length seemed to increase exponentially as vessel diameter decreased (**Supplementary Fig. 3c**).

The distance between pericyte cell bodies (inter-soma distance) also increases as vessel diameter decreased (Hall, C., personal communication).^28^ As a test of self-consistency for the cell length data, we returned to optically-cleared tissue from PDGFRβ-tdTomato mice to measure inter-soma distance of pericytes at locations relevant to ensheathing pericytes and mesh/thin-strand pericytes, *i.e.*, upstream and downstream of the α-SMA terminus (**Supplementary Fig. 6**). We collectively referred to mesh and thin-strand pericytes as capillary pericytes since they could not be distinguished with contiguous tdTomato labeling. Indeed, the inter-soma distance of ensheathing pericytes was significantly shorter compared to capillary pericytes.

Finally, we examined the extent of vessel coverage offered by each mural cell type. Coverage was calculated from average projected images as the percentage of tdTomato labeled area (**Fig. 5e-h**; red) divided by a larger area demarcated by lectin-labeled endothelium (**Fig. 5e-h**; black). Calculating tdTomato-labeled area involved applying a threshold for tdTomato intensity similarly across all mural cell groups. We chose a threshold with best sensitivity for differences between the three pericyte sub-types. This may have limited sensitivity to differences between smooth muscle cells and ensheathing pericytes. With this caveat in mind, vessel coverage between smooth muscle cells and ensheathing pericytes was not statistically different with our analysis method. However, coverage by ensheathing pericytes (>95%) was significantly higher than mesh pericytes (72%), and thin-strand pericytes (51%)(**Fig. 5j and Table 1**). We detected a trend toward higher coverage by mesh pericytes compared to thin-strand pericytes, but this did not reach statistical significance (**Fig. 5j**). In contrast to cell length, mural cell coverage decreased exponentially as vessel diameter decreased (**Supplementary Fig. 3d**). This range for mesh and thin-strand pericyte coverage is comparable to values reported in normal mouse cerebral vasculature (∼75%).^29^-^31^

## DISCUSSION

The goal of this study was to survey the diversity of mural cell types as arterioles branched into capillaries in the adult mouse cortex. We further set out to determine whether transitional forms of pericytes could be reliably distinguished from canonical smooth muscle cells and pericytes. We categorized all mural cells on penetrating arteriole offshoots (1^st^ order and downstream branches) as pericytes since all cells examined on these microvessels had protruding ovoid cell bodies, a feature historically used to define pericytes in brain and other organs. We further subcategorized the pericytes into three groups based on the appearance of their processes: Ensheathing, mesh and thin-strand pericytes, with the last being the canonical pericyte type and the former two being transitional forms. These pericytes occupied sequential territories on pre-capillary arterioles and capillaries. The extent of these territories depended upon the size of the penetrating arteriole offshoot.

Detailed comparisons of the characteristics of individual cells revealed that only ensheathing pericytes could be reliably distinguished from other mural cell types based on cell morphology, α-SMA content, and vascular location. In contrast, mesh and thin-strand pericytes overlapped heavily in the metrics we examined. We therefore conclude that mesh and thin-strand pericytes should be considered as one group, termed “capillary pericytes”, as they likely exist within a continuum. In contrast, ensheathing pericytes should be considered a transitional mural cell form distinct from smooth muscle cells and capillary pericytes.

### What is an ensheathing pericyte and a capillary pericyte?

These new data allow us to provide clearer definitions for pericyte types that adorn brain microvessels (**Fig. 6a**). Ensheathing pericytes are α-SMA-positive and occupy proximal branches of penetrating arteriole offshoots. Here, we define the pre-capillary arterioles as the microvessels occupied by ensheathing pericytes. Ensheathing pericytes cover the vessel nearly 100% based on our method of analysis, but differ from smooth muscle cells because they possess a protruding ovoid cell body and a more elongated shape. They also resided on 1^st^ to 4^th^ order branches, that averaged 9 μm in diameter rather than 16 μm for penetrating arterioles. In contrast, capillary pericytes, comprised of mesh and thin-strand pericytes, are α-SMA-negative, longer in total cell length, and bear processes of varying complexity that only partially cover the vessel, *i.e.*, 40-80%. Capillary pericytes occupy the capillaries, defined as microvessels occurring after the α-SMA terminus.

**Figure 6.**
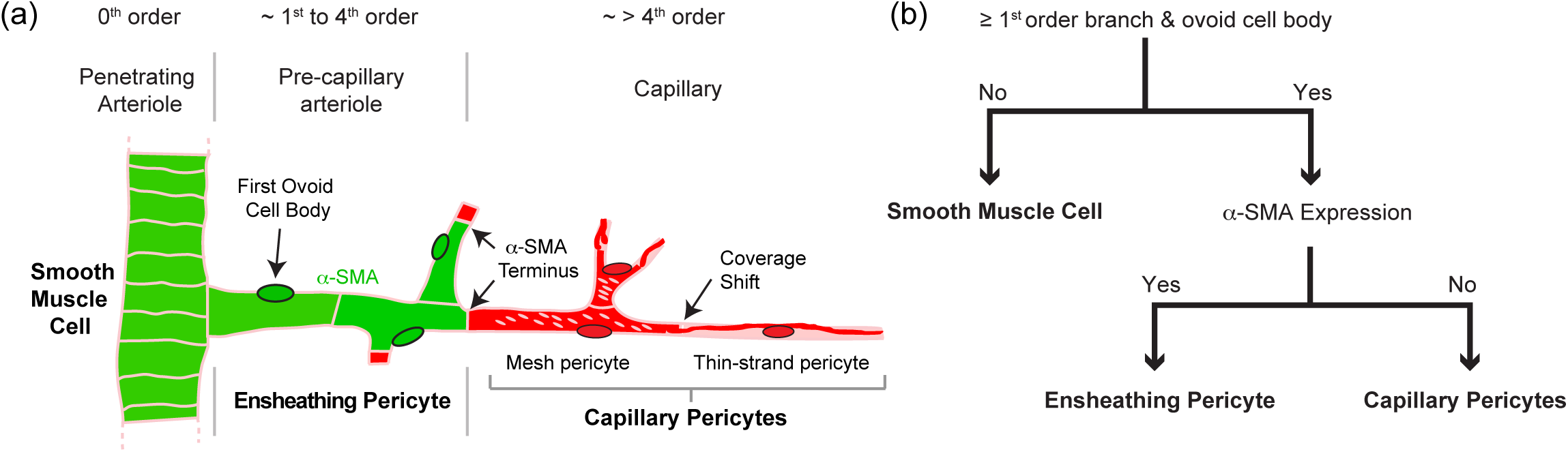
Organizational hierarchy for mural cell in mouse cortical vasculature and flow chart for classification. (**a**) Schematic showing transition of mural cell types as penetrating arteriole offshoots transition from pre-capillary arterioles to capillaries. The names of mural cell groups, features examined in this study, and approximate branch orders of occurrence are depicted. “Mesh pericytes” and “thin-strand pericytes” become more descriptive references to cell appearances within broader category called “capillary pericytes”, since they are difficult to separate using the metrics collected in this study. (**b**) Flowchart for distinguishing three major mural cells groups, smooth muscle cells, ensheathing pericytes, and capillary pericytes, the latter of which consists of mesh and thin-strand pericytes.

To further facilitate the classification of mural cells, we provide a flow chart (**Fig. 6b**) that relies only on knowledge of vascular topology and α-SMA expression. We anticipate that this flow chart and nomenclature will improve clarity in future discussions of cortical pericytes in normal and pathophysiological brain states. Oftentimes, there is insufficient information in studies to determine if results apply to ensheathing pericytes, capillary pericytes, or both, which likely have different contributions to blood flow regulation and other vascular functions.

### Identifying ensheathing pericytes and capillary pericytes in vivo

Studies seeking to differentiate between ensheathing and capillary pericytes on penetrating arteriole offshoots would benefit from an α-SMA label, such as that provided by the SMA-mCherry mouse line used by Hill *et al.*^13^ Our data further suggest that ensheathing pericytes reside only between 1^st^ to 4^th^ order branches. This information may be useful for conditions in which information on α-SMA expression is not attainable. For example, targeting branches beyond 4^th^ order ensures one is targeting capillary pericytes, while targeting branches under 4^th^ order, on large penetrating arteriole offshoots, increases the likelihood of studying ensheathing pericytes, but does not guarantee it. At the time of writing this article, Damisah *et al*. discovered that pericytes selectively uptake NeuroTrace 500/525 when topically applied to cortex *in vivo*, providing a fluorescent label for pericytes *in vivo*.^32^ The authors reported that the dye labeled only pericytes downstream of mural cells expressing α-SMA, suggesting that the dye was selective for capillary pericytes, and not ensheathing pericytes. We tested this dye in PDGFRβ-tdTomato mice and have reproduced the robust capillary pericyte labeling they demonstrated (**Supplementary Fig. 7**). Indeed, it appears that NeuroTrace 500/525 and α-SMA labeling are inversely related, but further quantification of NeuroTrace 500/525 intensity and α-SMA expression levels would help to confirm that these labels are truly mutually exclusive.

### Mural cell semantics

It should be noted that the cells described here as “ensheathing pericytes” have also been referred to as “pre-capillary smooth muscle cells” by other groups.^13, 14^ Indeed, ensheathing pericytes possess features of both pericytes and smooth muscle cells.^6^ In this study, we chose to subcategorize these cells as a form of pericyte for two reasons: 1) the protruding cell body has been a marker for pericytes across decades of research in many tissue types, and 2) α-SMA-positive pericytes have been described in past literature, though their vascular locations not well described^24^-^26^. Regardless of how to name this transitional cell type, which may be arbitrary until more information becomes available, it is important for future studies to describe the cortical mural cell types under investigation whenever possible. This will lead to a better understanding of the distinct roles of pericyte sub-types, and help identify potential sources of inconsistency.^7^

### Microvessel diameter and pericyte type

Ensheathing pericytes tended to occupy microvessels that were on average 3-4 μm larger (∼140%) than capillary pericytes. Yet, the diameters overlapped greatly between mural cell groups, and ensheathing pericytes could not be statistically separated from mesh pericytes based on diameter (**Supplementary Fig. 3b**). This suggests that pericytes could be either α-SMA-positive or α-SMA-negative if relying only on vessel diameter. Further, when interpreting the study by Hill *et al.*^13^ using the mural cell classification presented here, it appears that the basal tone applied by ensheathing pericytes *in vivo* can make the diameter of pre-capillary arterioles indistinguishable, and sometimes even smaller than nearby capillaries. Pre-capillary arteriole and capillary diameter can further change with the anesthetic state of the animal, or when considering *in vivo* versus *ex vivo* preparations.^33^ For these reasons, microvessel diameter alone is unlikely to be a reliable means to differentiate between ensheathing and capillary pericytes.

### Are ensheathing pericytes contractile in vivo?

While pericytes have been studied in retina, olfactory bulb, spinal cord and several peripheral organs, the vascular topology and organization of mural cells in these tissues may differ from arterioles of the cerebral cortex. We therefore have restricted our discussion of *in vivo* pericyte contractility to studies of the cerebral cortex.

Several studies have examined whether pericytes regulate blood flow in the intact brains of live mice using *in vivo* two-photon imaging. These studies appeared to report opposing results, with some groups suggesting a role for pericytes in physiological blood flow regulation^8^-^11^, while other groups suggested the contrary.^12^-^14^ Upon closer inspection, however, it becomes evident that there is consistency in the finding that capillary branches close to penetrating arterioles, *i.e.* 1^st^-4^th^ order branches, occupied by what we call ensheathing pericytes, exhibit vessel diameter change during normal brain activity. Hill *et al.* referred to these cells as pre-capillary smooth muscle cells and reported their spontaneous and optogenetically-induced contractility.^13^ Similarly, Fernandez-Klett *et al.* showed vasoreactivity at proximal branches with bicuculline-induced neuronal activation and spreading depression.^34^ Thus, when interpreting previous studies in light of the classification scheme presented here, several studies seem to agree that ensheathing pericytes regulate blood flow *in vivo*.

### Are capillary pericytes contractile in vivo?

Capillary pericytes are by far the most common pericyte form, and understanding their role in blood flow control is crucial. Past studies in brain slice by Attwell and colleagues have shown that about 50 to 70% of pericytes stimulated by agonists cause local capillary diameter change.^8, 35^ This suggests that a greater proportion of pericytes than just ensheathing pericyte have the capacity to contract. Indeed, *in vivo* two-photon imaging studies revealed that 25% to 30% of capillaries 4^th^ order or greater dilated in response to electrical stimulation of the limb.^8^ More recently, Kisler *et al.* also reported small dilations (1% on average) at cell bodies of capillary pericytes (thin-strand pericyte appearance in their presented images), and not on capillary regions devoid of pericytes.^10^ It was therefore surprising that Fernandez-Klett reported no changes of capillary diameter in response to seizure-like activity evoked by intracortical bicuculline injection.^12^ Wei *et al.* found no difference in basal capillary diameter or activity-evoked hyperemia when comparing regions covered and uncovered by capillary pericyte somata.^14^ Finally, direct optogenetic stimulation of capillary pericytes in the study by Hill *et al*. also did not yield any perceptible change in vessel diameter.^13^ Whether these discrepancies can be explained by differences such as imaging resolution, the animal’s physiological state, or strength and type of stimulation remains to be determined.

There is also evidence to suggest that capillary pericytes are involved in aberrant constriction during ischemia. Studies of capillary diameter during ischemia indicate capillary constriction that may be in part due to pericyte contraction.^9, 36^-^38^ While Hill *et al* suggest that the most prominent constriction occurs at the pre-capillary arteriole^13^, we have seen capillary constriction more broadly through the capillary bed.^36^ Are capillary pericytes involved in ischemic capillary constriction? Unpublished studies from our group^39^ suggest that capillary pericytes can indeed reduced capillary diameter and red blood cell flow *in vivo* with stronger optogenetic depolarization than that used by Hill *et al*.^13^ Cells with capillary pericyte morphology also contract during *in vivo* application of U46619, a TBXA2 receptor agonist and potent vasoconstrictor.^34^ Thus, there is evidence that strong depolarization or stimulation of capillary pericytes in pathological scenarios such as stroke may lead to sustained capillary constriction, such as with pericyte “rigor” and no-reflow.^8^ Actin isoforms other than α-SMA, *i.e.,* smooth muscle γ-actin, are reportedly expressed in pericytes^40^, and may be involved in pathological pericyte contraction.^41^

### Limitations and future steps

One limitation is that our study relied on histological measures of α-SMA, opening the possibility that we could not detect low levels of α-SMA. However, previous studies using α-SMA-CreER animals crossed with Cre reporters also showed no α-SMA in capillary pericytes.^13^ Further, recent single cell RNAseq studies^42, 43^ revealed that some mural cells had substantial α-SMA mRNA, while others had none.^38, 39^ Although our immunohistochemistry of α-SMA may not be as sensitive as transgenic labels of α-SMA or single-cell RNAseq data, it is important to consider that immunohistochemistry provides a direct measure of protein levels, whereas the other metrics rely on α-SMA promoter activity or RNA levels. Nevertheless, the preponderance of evidence using multiple methods suggests that capillary pericytes do not express α-SMA.

Another limitation is that our study focused on the arteriole pole of microvasculature due to its relevance in neurovascular coupling. Further, we focused on larger penetrating arterioles, and mural cell distribution may differ for smaller penetrating arterioles. Cells with transitional features also exist on post-capillary venules, and these cells may be involved in regulation of immune cell entry.^44^ We suspect that the approach used here will not provide clear classifications for venular pericytes, since α-SMA expression is generally lower and less clear-cut in venules.^1, 13^ It is also unclear if our approach to categorize mural cells in mouse cortex can translate to vasculature in other brain regions. While similar gradients in α-SMA expression and vessel coverage has been reported in other organs^26, 45^, pericyte density, morphology and vascular topology may differ greatly from brain^46^. Pericyte organization may also differ at various stages of brain development. In fact, a recent study found it difficult to distinguish smooth muscle cells and pericytes as α-SMA was not expressed in embryonic brain microvessels.^47^ Finally, it is important to note that pericytes are more diverse than the structural classification provided here, with roles in blood-brain barrier integrity^29, 30, 48^ and blood-brain barrier degradation during pathology^49, 50^, immune cell entry^44^, new vessel formation^51^, and potentially, the production of pluripotent stem cells.^23^ The results obtained from powerful genetic and biochemical approaches recently used to dissect pericyte genetics^42, 52^ need to be related to pericyte morphology and vascular location. This will help to link pericyte expression profiles with functional roles revealed by *in vivo* studies.

## Acknowledgements

We thank Catherine Hall and Pablo Blinder for helpful discussions. We also thank Ashley Watson, Manuel Levy and Christopher J. Guerin for critical reading of the manuscript.

## Author Contribution Statement

R.I.G., D.A.H., and A.Y.S. designed, executed and analyzed the studies. R.I.G. and D.A.H. wrote the manuscript with feedback from R.G.U., A.A.B., N.R.B and A.Y.S.

## Conflicts of interest

The authors have no financial or non-financial conflicts of interest.

Supplementary material for this paper can be found at http://jcbfm.sagepub.com/content/by/supplemental-data

